# CaMKII generates actin bundle morphology and mechanics distinct from canonical bivalent cross-linkers

**DOI:** 10.64898/2026.09.16.752118

**Authors:** Sofia Vargas-Hernandez, Carlos Bueno, Madiha Syeda, Joseph Pham, Lydia L. Mireles, Christina M. Di Mauro, Peter G Wolynes, M. Neal Waxham, Anna-Karin Gustavsson

**Author notes:** Division of Developmental Biology, Cincinnati Children’s Hospital Medical Center, Cincinnati, Ohio, USA.

## Abstract

Actin cross-linkers are essential modulators of the actin cytoskeleton, enabling structural diversity and dynamic remodeling. Among them, calcium/calmodulin-dependent protein kinase II (CaMKII) is unique in serving a dual role as a kinase and as a multivalent structural binder of actin in dendritic spines, where it plays a major role in supporting dendritic spine structure via its β subunit. To understand the basis of the structural function of CaMKII, we quantified the morphology and mechanics of actin bundles formed by CaMKII and directly compared them with bundles assembled by the canonical bivalent cross-linkers α-actinin and fascin. Using fluorescence microscopy, we measured contour length, straightness ratio, and persistence length across different cross-linker concentrations. Fascin generated progressively shorter, straighter, and stiffer bundles with increasing concentration, whereas α-actinin bundles remained largely unchanged. In contrast, increasing CaMKII concentration reduced both bundle straightness and persistence length, revealing a unique concentration-dependent increase in bundle flexibility. To investigate the structural origins of these mechanical behaviors, we complemented the experiments with coarse-grained simulations of cross-linked actin bundles. Simulations revealed that CaMKII-generated bundles retained a substantially larger fraction of their curvature in their time-averaged configuration than bundles formed by α-actinin or fascin, indicating that bundle mechanics are strongly influenced by organizational features beyond thermal bending fluctuations alone. CaMKII bundles displayed behavior consistent with additional structural complexity arising from their multivalent architecture and flexible linker domains. Together, our experimental and computational results identify CaMKII as a structurally and mechanically distinct actin cross-linker. We propose that the combination of multivalency and linker flexibility enables CaMKII to function as a molecular structural pivot, promoting flexible, adaptable actin assemblies that are mechanically compatible with the dynamic remodeling required for dendritic spine plasticity.

**Significance:** Actin cross-linkers play a central role in cytoskeletal architecture and mechanics, yet how multivalent cross-linkers differ from canonical bivalent proteins remains incompletely understood. CaMKII is unique in serving both as a kinase and a multivalent actin cross-linker in dendritic spines. By directly comparing CaMKII with fascin and α-actinin, we show that increasing CaMKII density uniquely reduces actin bundle stiffness, while simulations reveal enhanced curvature retention within CaMKII bundles. Together, these results suggest that CaMKII multivalency and flexible linker domains enable organizational modes and mechanical responses inaccessible to conventional bivalent cross-linkers. This work provides a biophysical basis for understanding how CaMKII supports the dynamic actin remodeling required for dendritic spine plasticity.

## Introduction

Actin is ubiquitous in cell biology, playing a critical role in cell function. The actin cytoskeleton is directly involved in general cellular processes such as migration, division, and mechanical support (1–3). It is also fundamental to specialized functions, such as structural plasticity in neurons, most extensively characterized in dendritic spines (4–9). Dendritic spines are small protrusive structures responsible for maintaining plasticity and connection between neurons (4, 5, 10). Actin polymers can perform such a myriad of functions because they interact with actin-binding proteins (ABPs). These proteins organize actin filaments into distinct assemblies with specific structural properties compatible with their function. Thus, to understand the role of the actin cytoskeleton in specific cellular functions, it is crucial to determine how specific ABPs modulate both the morphological and mechanical characteristics of the actin assemblies.

One major class of ABPs is cross-linkers, which connect individual actin filaments into higher-order structures. Two well-known examples of actin cross-linkers are fascin and α-actinin. Fascin is a small globular protein of ∼55 kDa (11) with physical size of around ∼6 nm (12). It cross-links actin in a parallel orientation, creating tightly packed bundles (13), which provides rigidity to protruding structures like filopodia (14). In contrast, α-actinin is a larger, rod-shaped homodimer with a molecular weight of ∼200 kDa (15, 16) and a length of ∼35 nm (17, 18). It cross-links mixed polarity filaments in loosely packed architectures (13, 19). Such structures are found in contractile and adhesive structures, such as stress fibers (18, 16).

While primarily known as a signaling enzyme, calcium/calmodulin-dependent protein kinase II (CaMKII) also has a structural function as an actin cross-linker in neurons (8, 9, 20–26). CaMKII is encoded by four genes that give rise to the α, β, γ, and δ isoforms, which exhibit distinct tissue distributions and cellular functions (27, 28). CaMKII subunits range in molecular weight between 54 - 72 kDa depending on the isoform (29, 30). Although CaMKII is highly enriched in the brain, where the α and β isoforms predominate, the γ and δ isoforms are more broadly expressed in non-neuronal tissues (25, 27). Among these isoforms, CaMKIIβ is the predominant actin-binding isoform in neurons and plays a central role in regulating the actin cytoskeleton and structural plasticity of dendritic spines (24, 31–33).

The enzyme possesses a unique structure consisting of an average of 12 subunits organized into a dodecameric double-decked central hub of ∼10 nm (34, 29, 35, 36). Disordered linkers connect each subunit within the hub to their catalytic domains that extend up to ∼36 nm (36, 37).

The actin-binding site in each CaMKIIβ subunit is located within this linker-catalytic region (22, 37, 38). Consequently, unlike canonical cross-linkers such as fascin or α-actinin, which are restricted to binding two filaments, CaMKII has a multivalency significantly higher than two (38, 39). While the role of CaMKII as an actin cross-linker is established (21, 23), it remains unclear how its multivalency and unique holoenzyme geometry influence the morphology and mechanics of the resulting actin networks. Quantitative characterization is therefore required to understand how this multivalent cross-linker modulates the structural rigidity of actin bundles. This knowledge is essential for gaining better insight into the properties of actin assemblies in highly specialized structures such as dendritic spines.

In this work, we characterize the morphological and mechanical properties of actin bundles cross-linked by fascin, α-actinin, and CaMKIIβ. From fluorescence images of these structures, we extract contour length, straightness ratio and persistence length. Contour length provides a measure of bundle extension, whereas the straightness ratio quantifies the overall geometric sinuosity of a bundle. Persistence length, by contrast, quantifies bending stiffness by measuring the correlation of a bundle’s direction over distance, and has been widely used to study cytoskeleton assemblies (40–43). Importantly, straightness ratio and persistence length capture distinct properties: straightness ratio describes the overall geometry of a bundle, whereas persistence length measures bending rigidity and reflects resistance to deformation. Together, these complementary metrics characterize both the architecture and mechanical properties of cross-linked actin bundles. Conducting these measurements side-by-side using an identical experimental setup and reaction conditions is crucial for properly contextualizing the biophysical properties of CaMKII bundles relative to the bundles produced from the canonical cross-linkers. We tested two concentrations of each cross-linker to assess how changes in protein density influence the mechanics of the resulting bundles. While such concentration titrations have been previously documented for fascin (40), to our knowledge, this has not been reported for either α-actinin or CaMKII.

Our work establishes CaMKII as a structurally and mechanically unique cross-linker. Our results show that increasing CaMKII concentration leads to a decrease in both bundle straightness and persistence length, a trend distinct from α-actinin, which exhibits stable morphology and mechanics independent of concentration, and from fascin, which forms stiffer, shorter filaments at higher concentrations. Furthermore, computational simulations provided novel results at length scales not accessible experimentally and revealed that CaMKII multivalency coupled with binding to actin via flexible linkers produces bundles with unique structural and mechanical properties relative to those formed with fascin and α-actinin. These findings provide a foundation for future studies of the structural role of CaMKII in dendritic spines and other actin-rich cellular compartments.

## Materials and methods

### Proteins

Rabbit skeletal muscle actin (>99% purity; AKL99-A, Cytoskeleton Inc.), wild-type human fascin 1 (>95% purity; CS-FSC01, Cytoskeleton Inc.), and rabbit skeletal muscle α-actinin (>85% purity; AT01, Cytoskeleton Inc.) were resuspended using general actin buffer (GAB) (1X buffer composition: 0.2 mM calcium chloride, 5 mM Tris, pH 8.0; BSA01-001, Cytoskeleton, Inc.). Aliquots were snap frozen in liquid nitrogen and stored at −80°C until use. The β isoform of CaMKII was used for all studies in this manuscript. Expression and purification of CaMKIIβ was accomplished using a baculovirus system exactly as described (23, 36, 44) and purification was carried out using calmodulin (CaM)-Sepharose affinity chromatography. Purified protein was dialyzed into 20 mM HEPES, 0.5 M NaCl, and 10% glycerol, pH 7.4 and stored as single use aliquots frozen at −80°C. CaMKII protein was quantified using a calculated extinction coefficient at 280 nm of 1.03 = 1 mg/ml. Using the calculated molecular weight of 60.4 kDa, this results in a protein concentration of 16.6 μM subunits, or 1.3 μM concentration of holoenzymes (25, 45).

### Sample preparation and electron microscopy (EM) imaging

F-actin was first polymerized from G-actin from rabbit skeletal muscle for 1 hr at room temperature as described by the manufacturer (Cytoskeleton Inc). Samples (25 μL reaction volume) for cryo-EM were prepared by diluting F-actin in actin polymerization buffer (APB) (1X strength APB composition: 10 mM Tris HCl, pH 7.5, 50 mM potassium chloride, 2 mM magnesium chloride, 1 mM ATP, 5 mM guanidine carbonate for ATP stabilization; BSA02-001, Cytoskeleton, Inc.) and separately adding the individual actin-binding proteins. The F-actin stock was diluted to a final concentration of 0.07 mg/ml actin, and then 0.04 mg/ml of fascin, α-actinin, or CaMKII were added, and incubated for an additional 30 min at room temperature with occasional mixing. The thawed aliquot of CaMKII was centrifuged at 14,000 rpm in an Eppendorf microfuge at 4°C to remove aggregates before addition to F-actin. For cryo-preservation, 5 μL of each sample was spotted onto a glow discharged Quantifoil grid (2/2), blotted from behind with Whatman 1 filter paper, and immediately plunge frozen in liquid ethane cooled to liquid nitrogen temperature. The grids were stored in liquid nitrogen until imaged. The grids were imaged at 200 kV using a FEI Polara F30 electron microscope equipped with a K2 Summit camera at a magnification 39,000x which gave a pixel size of 2.7 Å/pixel. Defocus was set to 5 to 10 μm and total electron dose was kept below 50 electrons/Å^2^. Thirty dose-fractionated frames were acquired for each image that were subsequently drift corrected using the MotionCorr2 package (46).

### Sample preparation for fluorescence imaging

Actin protein at 2.33 μM was incubated for 45 min at room temperature with 0.5 mM dithiothreitol (DTT) (R0861, Molecular Biology), 0.2 mM adenosine 5’ triphosphate disodium salt (ATP) (BSA04, Cytoskeleton Inc.), and 1X strength of APB. Polymerized actin was diluted to 1.5 μM in GAB and mixed with fascin, α-actinin or CaMKII at two different concentrations: 75 nM and 375 nM; resulting in molar ratios of 20:1 and 4:1 actin:cross-linker, respectively. These specific concentrations were selected based on previous fluorescence imaging studies of actin bundles (13, 40), with molar ratios chosen to ensure a significant difference in cross-linking density between the two regimes. The mixtures of cross-linkers and actin were gently mixed at room temperature using a tube revolver (88881001, Thermo Scientific) set to 40 rpm for 30 min. Next, phalloidin conjugated to Alexa Fluor 647 at 66 nM (A22287, Thermo Fisher) was added. Alexa Fluor 647-conjugated phalloidin was added after bundle formation to minimize the possibility that influences the cross-linking process itself. Then, the samples were mixed in the tube revolver for an additional 5 min. To remove large aggregates of CaMKII, the holoenzyme was centrifuged for 30 min at 14000 rpm at 4°C before mixing, and the supernatant was used to prepare the CaMKII-actin bundles.

For fluorescence imaging, the cross-linked actin bundles were plated on the glass bottom of an 8-well sample chamber (80827 Ibidi GmbH). The chamber was previously plasma cleaned with argon for 15 min and coated with poly-D-lysine (A-003-E, Thermo Fisher) diluted to 10 μg/mL in 1X phosphate buffered saline (PBS) (SH30256.01, Cytiva) to promote bundle adherence. Samples were kept moist by the addition of ∼150 μL of 1X PBS within 1 min following plating. To reduce photobleaching ∼150 μL of an oxygen-scavenger (GLOX) buffer was added to the PBS in the sample immediately before imaging. Briefly, to prepare the GLOX buffer, 14 mg glucose oxidase (GS2133, Sigma-Aldrich) and 50 μL catalase suspension (C100, Millipore Sigma) were added to 200 μL PBS. This solution was centrifuged for 5 min at 1000 x g and 4°C and the pellet was discarded. Then, 10 μL of this solution was combined with 690 μL nanopure water, 200 μL 50% (w/v) glucose (50-99-7, Sigma-Aldrich) and 100 μL 1M Tris-HCl (J22638.K2, ThermoFisher Scientific).

### Fluorescence microscopy data acquisition

Samples were imaged on an inverted microscope (IX-83, Olympus) using a 100X oil immersion objective (UPLAPO100XOHR, 100X, NA 1.5, Olympus), using a 647 nm laser (2RU-VFL-P-1000-647-FCAPC, 1 W, MPB Communications) with roughly matched illumination intensities of 3.2 mW/cm^2^ in homogeneous flat-field total internal reflection fluorescence (FF TIRF) illumination mode previously described (47). Between 10-20 tiff stacks of 10 frames of different fields-of-view (FOVs) were acquired at 50 ms exposure time using an sCMOS camera (Orca Fusion BT, Hamamatsu) with a calibrated pixel size of 130.8 nm. For each of the six conditions, three independent biological replicates were acquired across different days. The number of analyzed bundles per replicate varied between 34-117, and the total number of bundles per condition ranged between 204-301.

### Image postprocessing and morphological analysis

Following a similar postprocessing pipeline as previously described (48) and using the open-source software ImageJ (49), each 10-frame image stack was summed to improve signal-to-noise ratio (Fig. S1). Background subtraction was then performed using a rolling ball filter of 5-10 pixels, followed by 0.35-0.4 contrast enhancement and a final Gaussian blur filter with a 0.5-0.8 sigma (the standard deviation of the Gaussian function) range. These specific parameter ranges were determined by visual inspection to maximize background reduction while preserving the bundle signal.

The ImageJ plugin RidgeDetection (50) was used to detect the bundles and create a binary mask of the postprocessed image. Using a custom-built ImageJ macro, bundles were selected for analysis by retaining only those structures that had the 50% highest intensity in each FOV. This filtering process used an intensity threshold similar to that described in (40). The resulting masks were then skeletonized using the standard Skeletonize function in ImageJ. The skeletonized images were visually inspected to verify that the skeletons accurately traced the bundles, and traces were manually corrected where necessary.

Using a custom-made MATLAB script, the skeletonized bundle outlines were fitted to a smoothing spline with a smoothing factor of 0.1 (Fig. S1). This value for the smoothing factor was determined by visually confirming that the high-spatial frequency artifacts due to pixelation were mitigated, while simultaneously ensuring the smoothed traces captured the larger-scale outline of the bundles. Finally, the contour length (*L*), end-to-end distance (*d*), and straightness ratio (*S_r_*) of each smoothed trace were extracted. All samples after the initial characterization run were imaged and analyzed blind to reduce bias.

### Persistence length analysis

To determine the persistence length, *l_p_*, of the cross-linked actin bundles, we followed an approach similar to previous work (51) by fitting the two-dimensional cosine correlation to equation [1]:

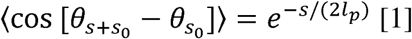

Using a custom-made MATLAB script, we extracted the differences between the tangent angle 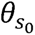 at a position *S*_0_ along the contour length and the corresponding tangent angle 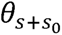 separated by a contour distance s. The distance *s* was incremented in 150 nm intervals. The maximum contour distance, *S_max_*, was set to be *L*/10, where *L* represents the total bundle length, as larger *s* values yield poor statistics (51). This resulted in a range 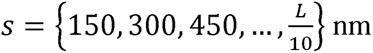.

For every given *s*, the cosine correlation values were averaged across each bundle to obtain 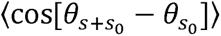. These averages were then fitted to equation [1] using a nonlinear least squares method. Bundles yielding fewer than three data points were excluded from the analysis. The quality of the non-linear fit was assessed by calculating the root mean square error (RMSE) for each bundle. The 95^th^ percentile RMSE remained below 0.03 across all conditions, indicating a consistently good quality of fit (Table S1).

### Statistical analysis

We calculated the median, first (Q1), and third (Q3) quartiles, as well as the interquartile range (IQR = Q3 - Q1), for the morphological parameters and the persistence length across each of the six bundle conditions. We selected these descriptive statistics because the data distribution is not symmetrical; thus, these measures provide a better representation of the data (Figs. S2, S3, S4).

**FIGURE 1.**
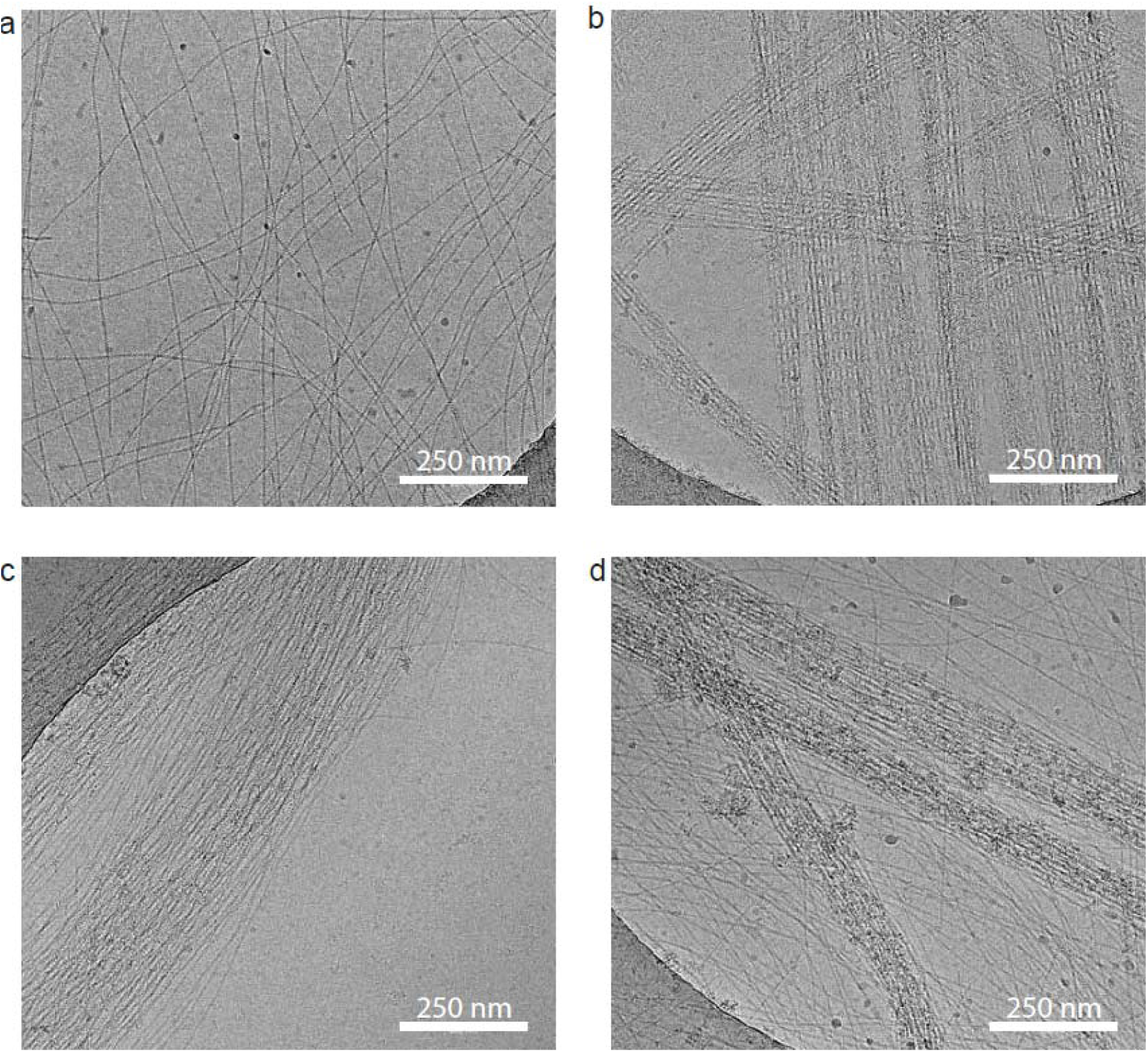
Cryo-EM images of actin filaments and actin bundles. Representative cryo-EM images of (a) individual actin filaments and actin bundles cross-linked by (b) fascin, (c) α-actinin, and (d) CaMKII. Note the random organization of actin filaments without cross linker (a) in contrast to the straight near crystalline organization of actin filaments when bundled by fascin (b). In comparison, α-actinin produces less densely packed curving bundles (c). CaMKII produces actin bundles intermediate in organization between fascin and α-actinin (d).

**FIGURE 2.**
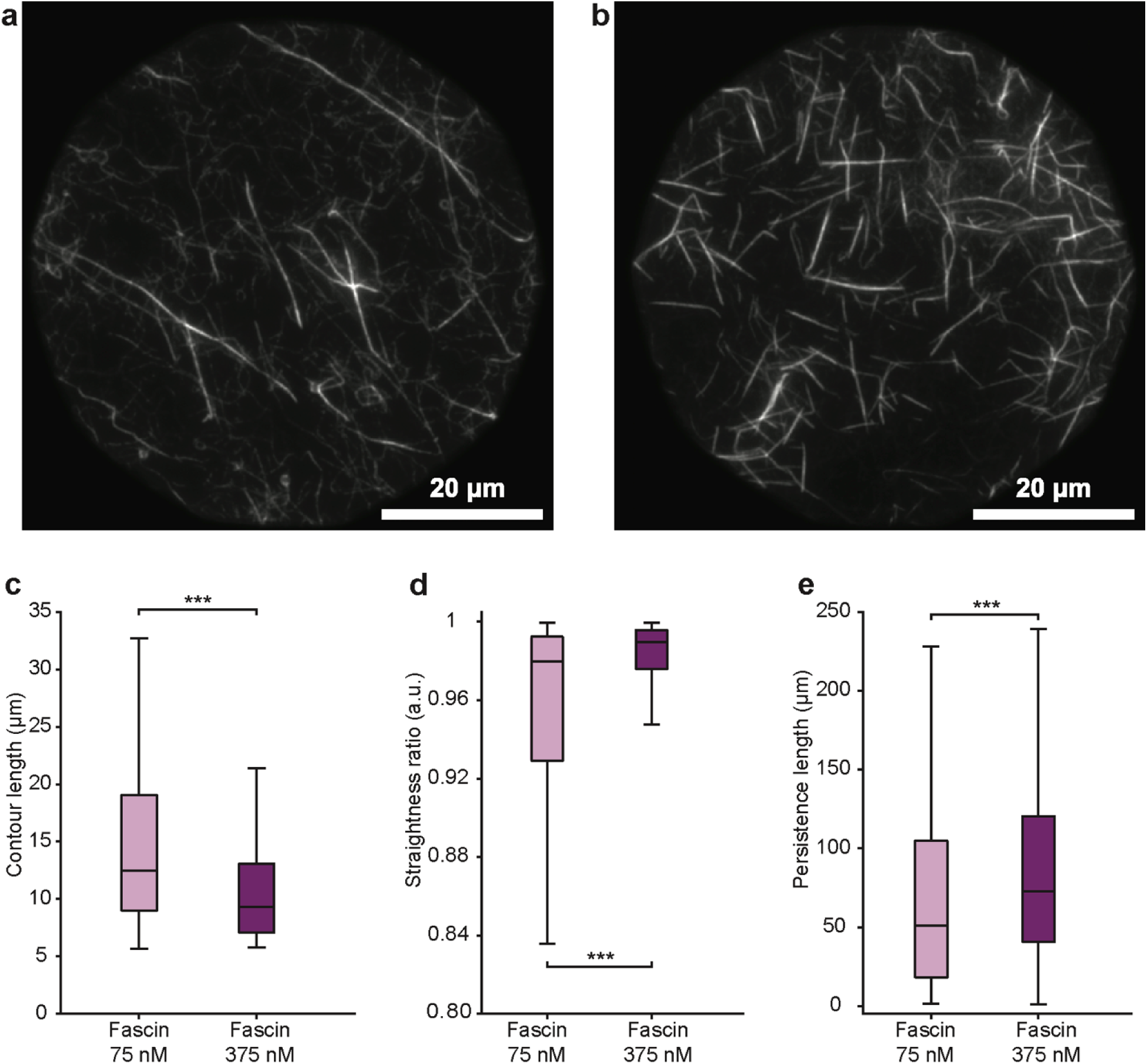
Increasing fascin concentration drives stiffening and limits bundle length. (a–b) Representative fluorescence images of actin bundles labeled with Alexa Fluor 647-phalloidin and cross-linked by fascin at (a) 75 nM and (b) 375 nM. Both images are normalized to the same contrast. (c–e) Box plots showing the bundle metrics from three independent replicates acquired across different days. Statistical significance was determined using a linear mixed model (LMM) to account for inter-replicate variability. (c) Contour length distributions showing a significant decrease in length at higher fascin concentrations (p < 0.001). (d) Straightness ratio measurements. Bundles are significantly straighter at high fascin concentrations (p < 0.001). (e) Persistence length calculations showing significant stiffening at higher concentrations (p = 0.004). Measurements were obtained from three independent blinded experimental replicates. The total number of bundles analyzed for each condition was 218 for fascin at 75 nM and 244 for fascin at 375 nM.

**FIGURE 3.**
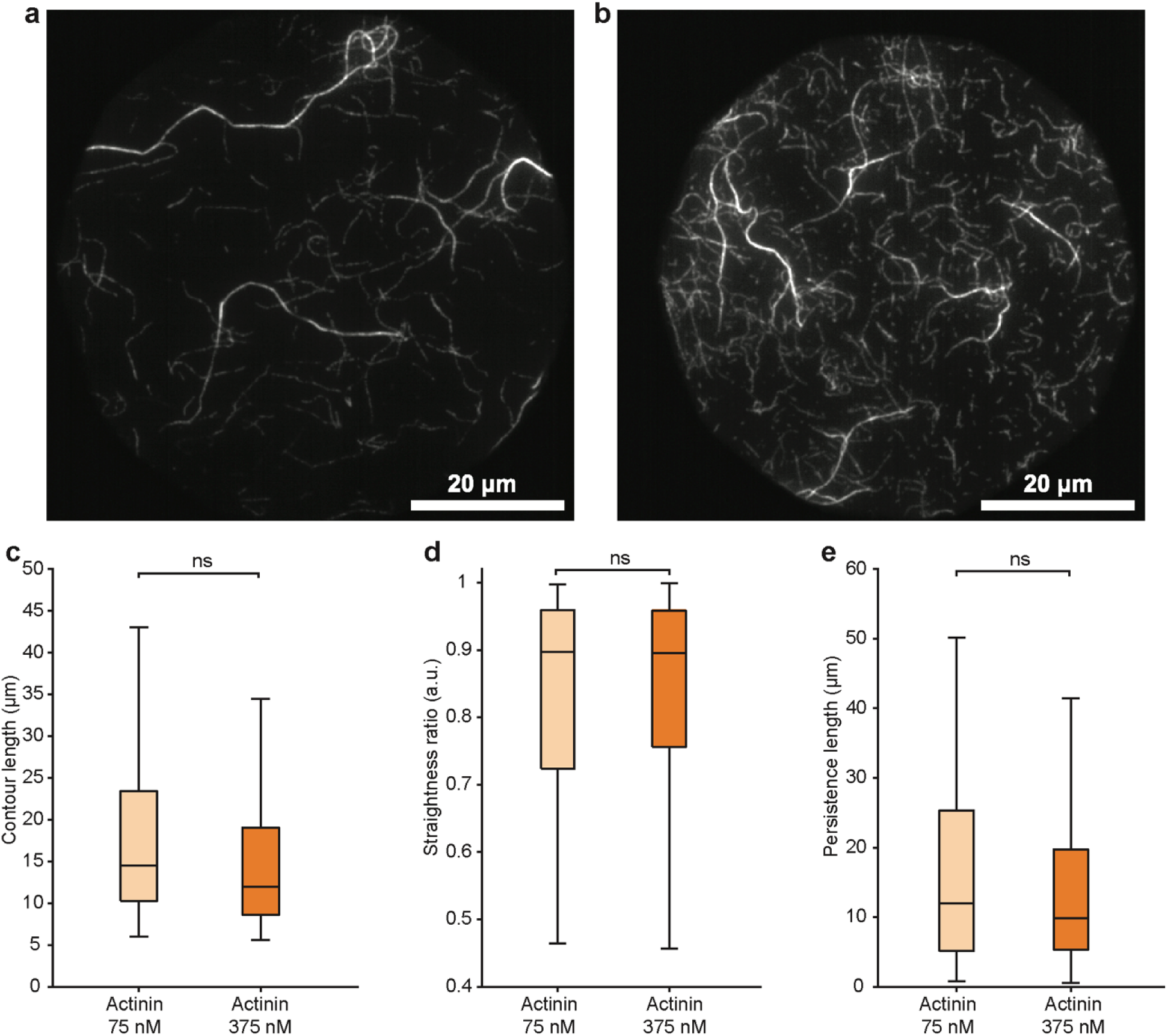
Bundle morphology and stiffness remain stable across α-actinin concentrations. (a–b) Representative fluorescence images of actin bundles labeled with Alexa Fluor 647-phalloidin and cross-linked by α-actinin at (a) 75 nM and (b) 375 nM. Both images are normalized to the same contrast. (c–e) Box plots showing the bundle metrics from three independent replicates acquired across different days. Statistical significance was determined using a linear mixed model (LMM) to account for inter-replicate variability. (c) Contour length distributions showing no significant changes across the two concentrations (p = 0.12). (d) Straightness ratio distributions showing no significant change with concentration (p = 0.98). (e) Persistence length calculations showing no significant change with concentration (p = 0.69). Measurements were obtained from three independent blinded experimental replicates. The total number of bundles analyzed for each condition was 183 for α-actinin at 75 nM and 222 for α-actinin at 375 nM.

**FIGURE 4.**
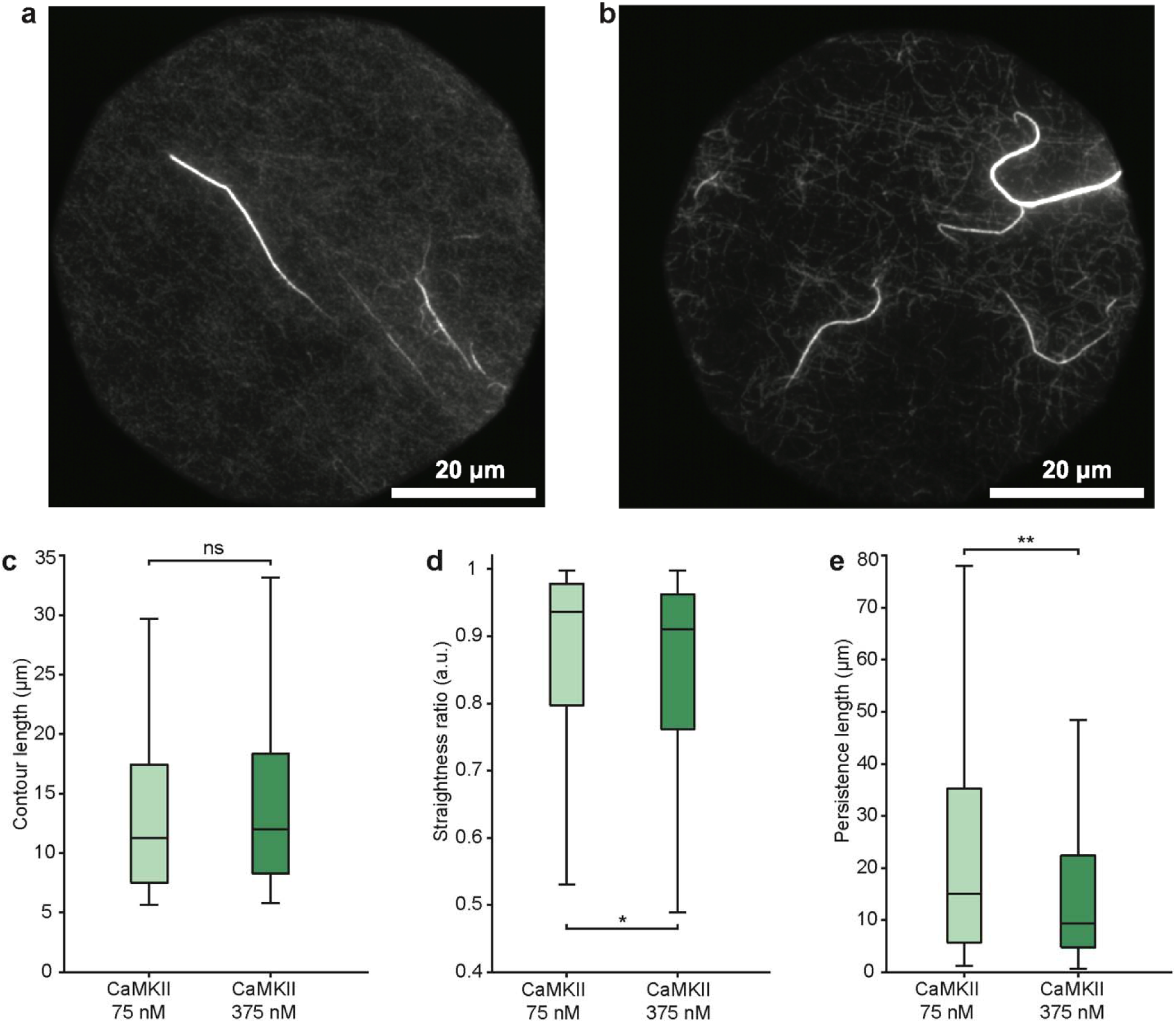
Higher CaMKII concentrations reduce bundle straightness and stiffness. (a–b) Representative fluorescence images of actin bundles labeled with Alexa Fluor 647-phalloidin and cross-linked by CaMKIIβ at (a) 75 nM and (b) 375 nM. Both images are normalized to the same contrast. (c–e) Box plots showing the bundle metrics from three independent replicates acquired across different days. Statistical significance was determined using a linear mixed model (LMM) to account for inter-replicate variability. (c) Contour length distributions showing no significant changes across the two concentrations (p = 0.19). (d) Straightness ratio distributions showing a statistically significant decrease with concentration (p = 0.023). (e) Persistence length calculations showing a significant decrease with concentration (p = 0.003). Measurements were obtained from three independent blinded experimental replicates. The total number of bundles analyzed for each condition was 162 for CaMKII at 75 nM and 194 for CaMKII at 375 nM.

To account for variability across the three independent replicates per condition (52–54), we analyzed the data using a linear mixed-effects model (LMM) using a custom MATLAB script. Prior to analysis, contour length, end-to-end distance, and persistence length were log-transformed, while the straightness ratio was logit-transformed to satisfy normality assumptions. Pairwise comparisons were performed between the two different concentrations for each parameter and each cross-linker, and also for different cross-linkers at the same concentration. For the latter comparisons, the p-values were adjusted for multiple comparisons using the Benjamini-Hochberg correction at a significance level of 0.05. This method was selected to maintain statistical power while controlling the proportion of false discoveries (55–57).

### Computational simulations of persistence length

Actin filaments were represented with the four-site coarse-grained model (58), one bead per structural subdomain of the protomer. Since actin subunits polymerize with ATP bound and hydrolyze it soon after incorporation, we used a model with the set of parameters corresponding to the actin-ADP bound state, matching mature filaments found in the experiments. All three cross-linkers were built from the same rigid three-bead actin-binding module and differ in how two or more modules (actin-binding sites) are joined. Fascin joins two modules by nine harmonic bonds that fix both the separation and the relative orientation of the bound filaments, with its two inequivalent binding faces taken from the cryo-EM structure of fascin cross-linked F-actin (PDB 8VO7, (59)). α-actinin joins two modules by a single 35 nm bond representing the spectrin rod. CaMKII carries twelve independent flexible modules on a rigid hub, with the hub and binding geometry taken from the structural model of the CaMKIIβ–F-actin complex (38) so that a single holoenzyme can engage more than two filaments simultaneously. For all crosslinkers each module binds through a smooth, reversible three-point potential well of depth ε, so that cross-links break and re-form over the course of a simulation.

Seven-filament hexagonal bundles of 416 monomers per actin filament and 1.1 µm of contour length were simulated in OpenMM 8 (60) with a Langevin integrator at 300 K, in eight independent replicas per condition. No explicit electrostatic term was included, so that cross-linker binding provides the only attractive interaction. The well depth ε was varied between 25 and 100 kJ/mol to represent differences in binding affinity, over the range in which each cross-linker holds a bundle together. Each trajectory was truncated to a common 10 µs, of which the first half was discarded as equilibration and the remaining 5 µs were analyzed.

Filament centerlines were extracted per frame as the centroid of the core beads of each monomer and averaged over one helical crossover. Before the mode amplitudes were computed, each configuration was rigidly aligned to the average structure. The bundle centerline was taken as the mean over the filament centerlines. Its unit tangent field was resolved into two orthogonal directions transverse to the mean bundle axis. Each of the transverse components was expanded independently in a series of cosine bending modes, indexed by, using the basis 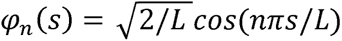 with L being the contour length. That gives us two amplitudes for each bending mode, 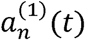 and 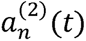, corresponding to the bending in the two directions and equation [2]:

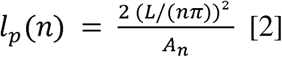

For each mode (*n*), the two transverse amplitudes (*a_n_*) were combined to obtain a single mode amplitude (*A_n_*) in two ways. In Equation [3], we used the variance of each amplitude about its time average to calculate the fluctuation persistence length, which measures bending fluctuations around the time-averaged bundle shape. In Equation [4], we used the mean square amplitude without subtracting that average to calculate the apparent persistence length (61, 62). Since the mean square is the variance plus the square of the mean, the two differ by the time-averaged shape alone.

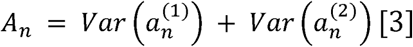

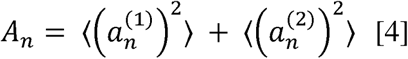

The cosine correlation of equation [1] was also fitted per bundle to the simulated centerlines. Since the simulated centerline is known in three dimensions, we fitted the three-dimensional form of equation [1], equation [5], which drops the projection and the factor of two in the exponent of equation [1].

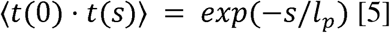

The cosine correlation was evaluated over separations from 100 nm to the end of the contour, at the spacing of one helical crossover, or 13 monomers.

Every persistence length was measured on each bundle separately and is reported as the median across the eight independent bundles of a condition, with the interquartile range as its uncertainty. Within each bundle the reported mode-based persistence length is the median of the mode-specific estimates for the bending modes from 4 to 10. Modes 1 to 3 were excluded since they were the least converged. Force field parameters and the protocol are detailed in the Supplemental Information. The parametrization of this force field will be described in a future article. The simulation code is available at https://github.com/cabb99/OpenActin/tree/persistence_length.

## Results and discussion

### Cryo-EM visualization of actin filaments cross-linked by fascin, α-actinin, and CaMKII

To corroborate the cross-linking capabilities of our fascin, α-actinin, and CaMKII proteins, we acquired cryo-EM images of individual actin filaments (Fig. 1 a) and cross-linked bundles (Fig. 1 b-d). These images confirm that all three proteins induce the formation of higher-order actin structures distinct from individual filaments and validate the biological activity of the preparations used throughout the study. The observed bundle morphologies are also consistent with previous EM characterizations (13, 19, 23, 63). Specifically, fascin generates tightly packed, straight, parallel bundles (Fig. 1 b). In contrast, α-actinin and CaMKII form looser assemblies that do not exhibit the highly periodic, crystalline-like packing characteristics of fascin bundles (Fig. 1 c,d). Having established that each cross linker generates its expected bundle architecture, we proceeded to quantify how each cross-linker modulates actin bundle morphology and mechanics.

### Increasing fascin concentration drives stiffening and limits bundle length

Fascin is well-known for creating actin bundles that provide structural support and are highly sensitive to cross-linker concentration (63). In our work, this sensitivity was reflected in several ways. First, increasing the concentration of fascin resulted in a significant decrease in the length of the imaged bundles (Fig. 2 a,b). The median contour length dropped by roughly 33%, decreasing from 11.7 µm (IQR = 8.1–18.5 µm) at 75 nM to 7.8 µm (IQR = 5.8–11.8 µm) at 375 nM (Fig. 2 c). The distribution of length values also narrowed at higher concentrations (Fig. 2 a,b), suggesting a homogenization of bundle lengths.

Fascin bundles also became significantly straighter at higher concentrations (Fig. 2 d). The median straightness ratio increased from 0.983 (IQR = 0.938–0.993) to 0.991 (IQR = 0.982– 0.996). As expected, this straightening was accompanied by a significant increase in persistence length (Fig. 2 e). The median persistence length increased by ∼35%, from 48.0 µm (IQR = 22.9– 78.1 µm) at 75 nM to 64.6 µm (IQR = 45.9–105.4 µm) at 375 nM. Although persistence lengths for fascin-cross-linked bundles at similar molar ratios have been reported previously, the absolute values vary substantially across studies, ranging from ∼30 µm to >100 µm depending on experimental designs (e.g., phalloidin stabilization), and analysis methods (40–42). Despite this variability in absolute values, the concentration-dependent stiffening observed here is consistent with previous work (40, 64). The consistency of this trend across studies, despite differences in the reported absolute values, highlights the importance of performing side-by-side measurements under identical experimental conditions, allowing biologically meaningful concentration-dependent responses to be distinguished from variations arising from experimental design and analysis methods. Furthermore, this mechanical reinforcement is consistent with the functional role of fascin, which provides the necessary rigidity for protruding cellular structures.

Mechanistically, this behavior reflects the geometry of the fascin cross-link. Because fascin is a small, globular protein that creates a tight, rigid interface, it accumulates structural strain within the bundle as it grows (65). It has been recently established that this strain, which arises from forcing actin filaments into a hexagonal lattice, thermodynamically limits bundle thickness (59). In our experiments, this accumulated strain likely contributes to the observed increase in stiffness but may also constrain longitudinal growth or, alternatively, increase the brittleness of the bundles. In the latter case, the shorter lengths observed at high concentrations may result from increased fracturing of these highly strained, rigid structures under the shear forces of pipetting and sample preparation.

### Bundle morphology and stiffness remain stable across α-actinin concentrations

In contrast to fascin, increasing the concentration of α-actinin did not produce significant changes in bundle morphology or persistence length of α-actinin-cross-linked bundles (Fig. 3). Even though there was a slight decrease in the median and variance of the contour length values, from 13.7 µm (IQR = 8.8–22.3 µm) at 75 nM to 10.6 µm (IQR = 7.5–17.2 µm) at 375 nM, the difference did not reach statistical significance (Fig. 3 c).

The straightness ratio also remained stable, with the median value and overall variance being 0.904 (IQR = 0.786–0.963) and 0.908 (IQR = 0.792–0.962) at 75 nM and 375 nM concentration, respectively (Fig. 3 d). Similarly, the persistence length was unaffected by concentration, with a median value of 16.3 µm (IQR = 8.5–28.4 µm) and 15.0 µm (IQR = 8.7– 27.1 µm) at 75 nM and 375 nM, respectively (Fig. 3 e). Importantly, the persistence length of these bundles is significantly lower than the median values observed for fascin. This trend aligns with previous mechanical characterizations (41, 42), which consistently identify α-actinin bundles as more flexible than fascin-cross-linked bundles. This is also evident in the cryo-EM images of the bundles (Fig. 1).

While previous studies noted the stability of α-actinin-cross-linked actin bundles under varying crowding conditions (41), the effect of varying α-actinin concentration on bundle mechanics has not, to our knowledge, been systematically examined. The rod-shaped α-actinin (18) antiparallel dimer acts as a rigid inter-filament spacer (∼35 nm) (13). Unlike the tight coupling of fascin, this wide spacing permit greater structural flexibility. Furthermore, once the bundle geometry is established, adding higher densities of this fixed-length spacer does not tighten the inter-filament coupling or induce additional stiffening, resulting in the mechanically stable regime observed in our data. This flexibility, combined with insensitivity to concentration changes, is biologically consistent with the role of α-actinin in actomyosin networks, since the wide spacing preserves the porosity required for molecular motors, such as myosin, to penetrate and contract the scaffold.

### Higher CaMKII concentrations reduce bundle straightness and stiffness

Increasing the concentration of CaMKII did not significantly change the contour length of the bundles, which maintained median values of 10.1 µm (IQR = 6.8–15.0 µm) and 10.9 µm (IQR = 7.3–17.7 µm) at 75 nM and 375 nM, respectively (Fig. 4 a-c). Strikingly, however, the bundle geometry behaved in direct contrast to both fascin and α-actinin. The straightness ratio significantly decreased with higher cross-linker concentrations from a median of 0.945 (IQR = 0.814–0.980) at 75 nM to 0.915 (IQR = 0.770–0.964) at 375 nM (Fig. 4 d).

This reduction in straightness ratio was consistent with a significant decrease in persistence length, which decreased by ∼33% at high concentrations; a magnitude of change comparable to that of fascin, but in the opposite direction. The median persistence length was reduced from 20.6 µm (IQR = 8.7–37.4 µm) at 75 nM to 13.8 µm (IQR = 7.4–24.9 µm) at 375 nM (Fig. 4 e). To our knowledge, this is the first report demonstrating that CaMKII concentration inversely regulates actin bundle stiffness.

One possible explanation for the observed reduction in persistence length is the unique architecture of the CaMKII holoenzyme acting as a molecular structural pivot. We define a ‘molecular structural pivot’ as a multivalent cross-linker that possesses sufficient internal conformational freedom, arising from its multivalency combined with its flexible linkers, to partially decouple the orientation of bound filaments from the geometry of the cross-linker. Under this framework, increasing CaMKII density may allow the bundle to adopt more sinuous configurations rather than progressively stiffening into a rigid lattice while maintaining excellent cross-linking stability.

### Primary mechanical distinction separates fascin from CaMKII and α-actinin

Next, we compared the morphological characteristics of these bundles for different cross-linkers at fixed concentrations. At 75 nM, bundle contour lengths are similar between fascin and α-actinin (median: 11.7 μm and 13.7 μm, respectively; p = 0.053), while CaMKII produced slightly shorter bundles (10.1 μm; p < 0.001 vs. α-actinin; p = 0.027 vs. fascin) (Fig. 5 a). However, at 375 nM, fascin bundles are significantly shorter (median 7.8 μm; p = 0.006 vs. CaMKII; p = 0.018 vs. α-actinin) than those formed by either α-actinin or CaMKII (median 10.6 μm and 10.9 μm, respectively) (Fig. 5 b), whereas the contour lengths of α-actinin and CaMKII are comparable (p = 0.52).

**FIGURE 5.**
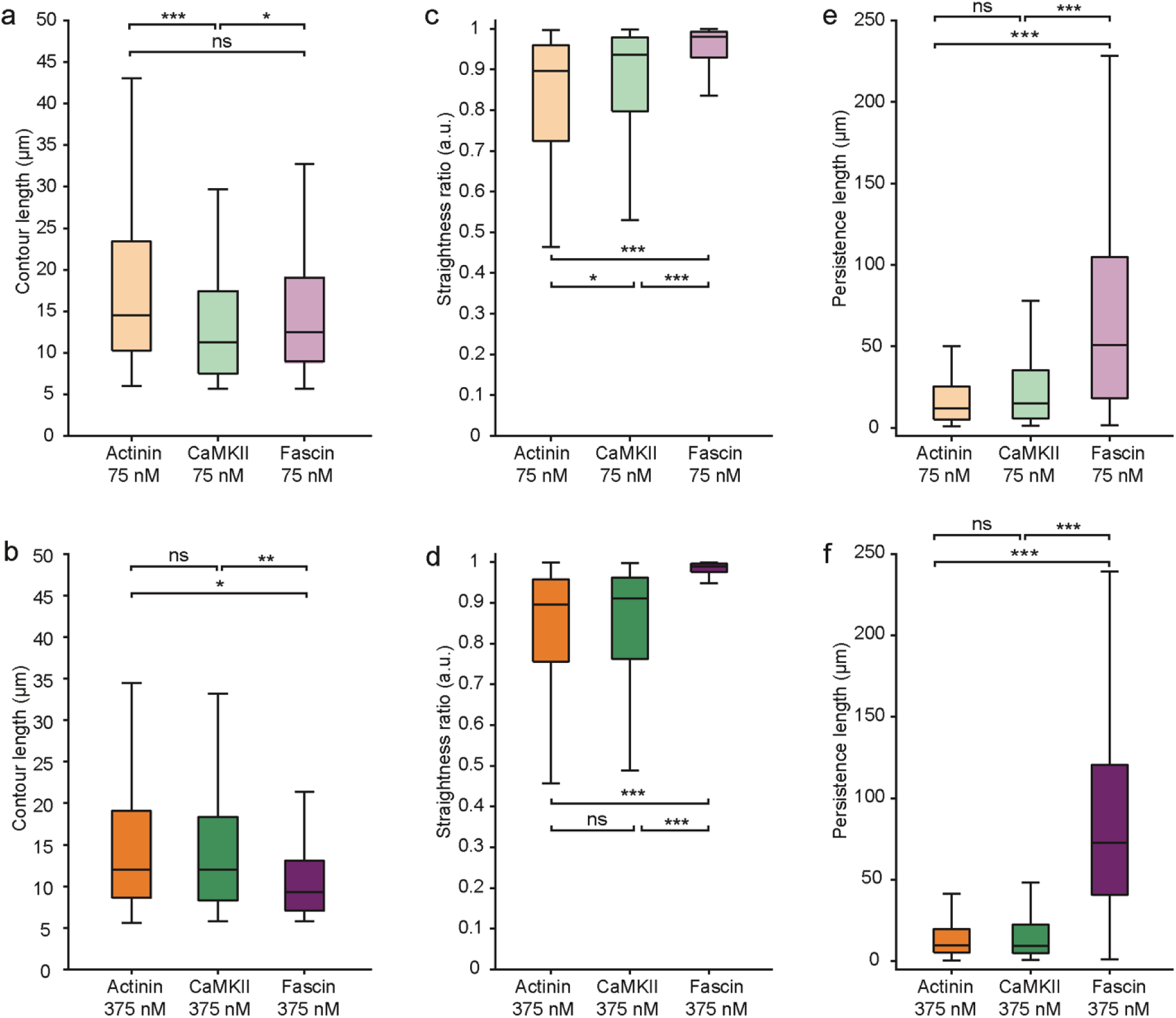
α-actinin and CaMKII form a distinct category of cross-linkers compared to fascin. (a–f) Comparisons of (a,b) contour length, (c,d) straightness ratio, and (e,f) persistence length distributions for actin bundles cross-linked by α-actinin, CaMKII, or fascin at (a,c,e) 75 nM and (b,d,f) 375 nM. The box plots show the bundle metrics from three independent replicates acquired across different days. Statistical significance was determined using a linear mixed model (LMM) to account for inter-replicate variability. p-values were adjusted using the Benjamini-Hochberg correction for multiple pairwise comparisons. (a) At 75 nM, the contour length differs significantly between α-actinin and CaMKII (p < 0.001) and fascin and CaMKII (p = 0.027), but there is no significant difference between α-actinin and fascin (p = 0.053). (b) At 375 nM, the contour length does not differ between α-actinin and CaMKII (p = 0.52), but significant differences are found between fascin and CaMKII (p = 0.006) and α-actinin and fascin (p = 0.018). (c) At 75 nM, the straightness ratios differ significantly across all proteins: α-actinin versus CaMKII (p = 0.018), fascin versus CaMKII (p < 0.001), and α-actinin versus fascin (p < 0.001). (d) At 375 nM, the straightness ratios differ significantly between fascin and α-actinin (p < 0.001) and fascin and CaMKII (p < 0.001), but not between α-actinin and CaMKII (p = 0.94). (e) At 75 nM, the persistence length differs significantly between fascin and α-actinin (p < 0.001) and fascin and CaMKII (p < 0.001), but not between α-actinin and CaMKII (p = 0.40). (f) At 375 nM, the persistence lengths differ significantly between fascin and α-actinin (p < 0.001) and fascin and CaMKII (p < 0.001), but not between α-actinin and CaMKII (p = 0.63). Measurements were obtained from three independent blinded experimental replicates. The total number of bundles analyzed for each condition was 218 for fascin at 75 nM, 244 for fascin at 375 nM, 183 for α-actinin at 75 nM, 222 for α-actinin at 375 nM, 162 for CaMKII at 75 nM, and 194 for CaMKII at 375 nM.

In terms of straightness ratio and persistence length, fascin formed significantly straighter and stiffer bundles (p < 0.0001) than the other two cross-linkers. At 75 nM, fascin bundles exhibited a median straightness ratio of 0.983 and persistence length of 48.0 μm; at 375 nM, these values increased to 0.991 and 64.6 μm, respectively (Fig. 5 c–f). At 75 nM, CaMKII produced slightly straighter filaments than α-actinin (median 0.945 vs. 0.904; p = 0.018). Interestingly, statistical analysis revealed no significant difference in persistence length between α-actinin and CaMKII at 75 nM (median 16.3 μm vs. 20.6 μm; p = 0.40) or at 375 nM (15.0 μm vs. 13.8 μm; p = 0.63), or in straightness ratio at 375 nM (median 0.908 vs. 0.915; p = 0.94). This indicates that both α-actinin and CaMKII create structures with similar overall sinuosity and flexibility (Fig. 5 e,f).

This similarity presents an interesting nuance. While our data shows that CaMKII tunes bundle stiffness with concentration, unlike α-actinin, which remains stable, this behavior does not shift it into the “rigid” category. Although CaMKII exhibits a significant ∼33% increase in flexibility (compliance) as concentration increases, the absolute values remain much closer to α-actinin than to fascin. For example, at 375 nM, the median persistence lengths of CaMKII (13.8 µm) and α-actinin (15.0 µm) are comparable, whereas fascin forms significantly stiffer bundles (64.6 µm) (Fig. 5 f). Therefore, even though CaMKII has a distinct dynamic response to concentration, the magnitude of the resulting bundles straightness and stiffness keeps it mechanically more similar to α-actinin than to fascin.

Although there are significant differences in morphology and mechanical response between CaMKII and α-actinin (Figs. 2, 3, and 5 a,c), our persistence length data indicate that the most significant mechanical difference lies between fascin and the other two cross-linkers. The similar magnitude of persistence length for CaMKII and α-actinin suggests a high degree of compatibility between these two proteins. This flexibility may facilitate the formation of actin architectures ranging from bundles to networks in dendritic spines (66). Thus, CaMKII may provide a flexible cross-linking mechanism capable of accommodating changes in actin organization while remaining mechanically compatible with α-actinin-rich assemblies. Under this interpretation, the distinct concentration-dependent behavior of CaMKII could allow actin structures to adapt to variations in cross-linker composition while maintaining mechanical flexibility.

While the physiological stoichiometries of these cross-linkers remain incompletely defined (67, 68), particularly within specific actin structures, the concentrations used here enable direct side-by-side comparison of their mechanical effects on actin bundles. Under these controlled experimental conditions, fascin, α-actinin, and CaMKII exhibit distinct behaviors, revealing fundamental differences in how these proteins modulate actin cross-linking and providing insight into how their distinct biophysical properties may contribute to their biological functions.

### Simulated CaMKII-cross-linked bundles retain a larger fraction of their curvature than fascin- or α-actinin-cross-linked bundles

To determine whether the distinct behavior of CaMKII reflects differences in bundle organization rather than simply differences in bulk stiffness, we performed computational simulations of actin bundles cross-linked by fascin, α-actinin, and CaMKII of 1.1 µm contour length (Fig. 6). These simulations provide access to length scales that are not experimentally accessible from our fluorescence measurements. To distinguish thermal fluctuations from curvature retained in the average bundle shape, we quantified bundle mechanics using fluctuation persistence length, apparent persistence length, and persistence length derived from cosine correlations. The fluctuation persistence length isolates the contribution of thermal bending fluctuations, whereas the apparent and cosine correlation persistence lengths are computed from bundle centerlines and therefore capture contributions from both thermal fluctuations and the average bundle shape, making them more directly comparable to the persistence lengths measured experimentally.

**FIGURE 6.**
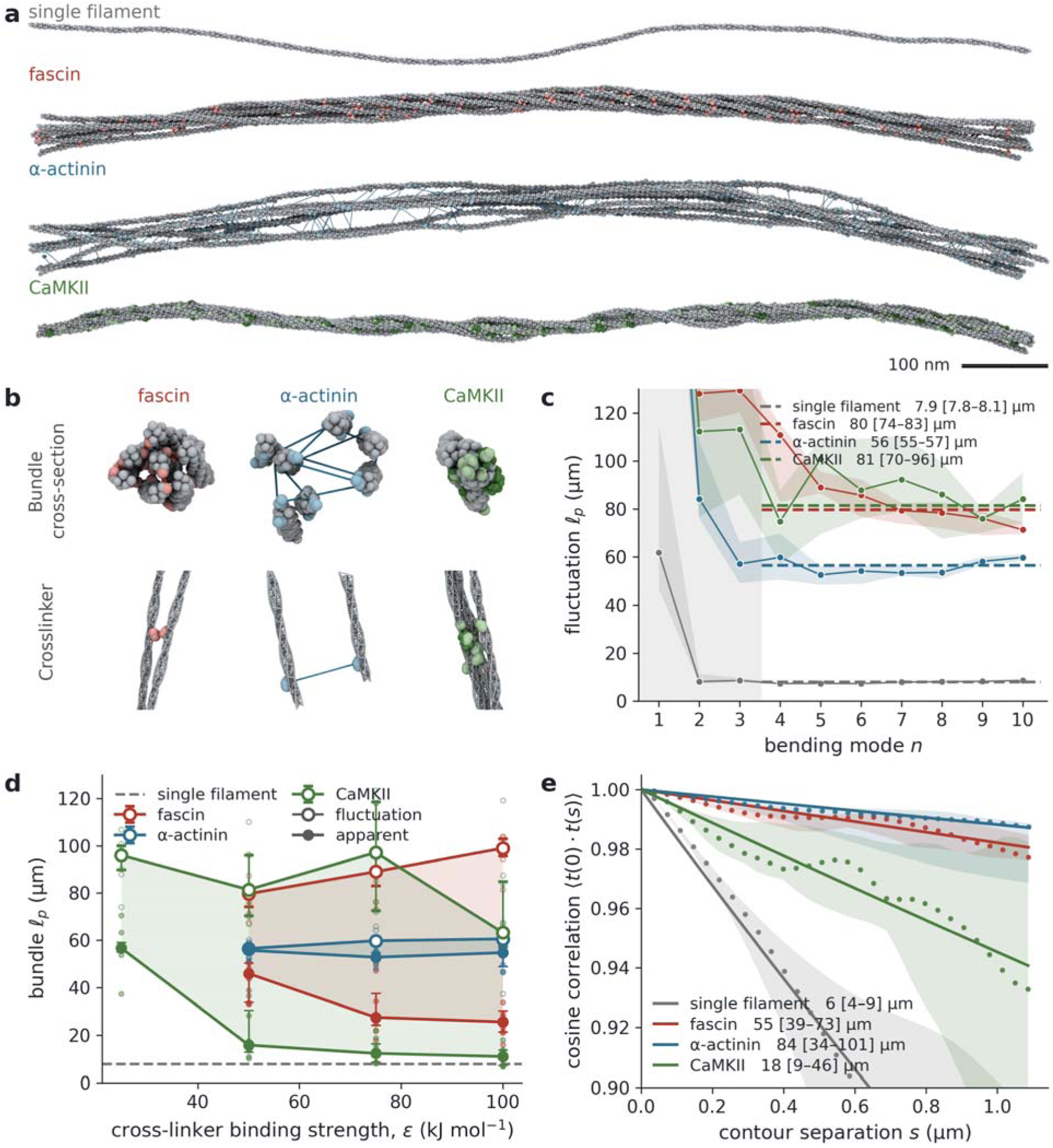
The apparent and fluctuation persistence lengths differ most in CaMKII bundles. (a) Side views of a single actin filament and representative bundles for fascin, α-actinin, and CaMKII. (b) Cross-sections of the simulated seven-filament bundles cross-linked by fascin, α-actinin, and CaMKII, cut perpendicular to the bundle axis, with cross-linker molecules shown within the slab. Shown below each cross-section is a single cross-linker with the filaments it binds, two for fascin, two for α-actinin, and four for CaMKII. Actin is grey and the cross-linkers are colored red, blue, and green for fascin, α-actinin, and CaMKII, respectively, with their actin-binding domains shown in a lightened color. The same colors are used in all panels. (c) Fluctuation persistence length. Points are the median over the eight replicates at each mode and the band shows their interquartile range, while the bold, dashed line is the plateau, calculated as the median persistence length over modes 4 to 10, given in the legend with their respective interquartile range. Modes 1 to 3 are greyed and excluded from the plateau calculation. (d) The fluctuation and apparent persistence lengths as a function of the well depth ε, drawn open and filled, respectively, where faint points represent the values for individual replicates. The shaded band is the difference between the two. The dashed grey line is a single actin filament (independent of ε). (e) Persistence length from the three-dimensional cosine correlation of the bundle centerline. Points are the median correlation over the eight replicates and the band their interquartile range, while the bold line is the exponential for the median of the eight fitted decay lengths, given in the legend with their respective interquartile range.

The persistence length of a single actin filament was 7.9 µm (IQR = 7.8–8.1 µm) from the bending-mode spectrum and 6.1 µm (IQR = 3.9–9.2 µm) from the cosine correlation, slightly below the 9.0 ± 0.5 µm reported for unstabilized actin filaments (69, 70), and 8.7 µm reported for this model in the ADP state (58).

The fascin and α-actinin models carry two binding modules and can hold at most two filaments. The valency of CaMKII is not imposed in this way, and the holoenzymes were found to hold generally two to five of the seven available filaments (Fig. 6 a,b). Individual CaMKII holoenzymes have been shown to cross-link multiple filaments (39), and the multidomain binding geometry along F-actin from which the model is built has been resolved (38), but the number of filaments one holoenzyme holds has not been experimentally measured.

At a binding energy of ε = 50 kJ/mol the fluctuation persistence length is 80 µm (IQR = 74–83 µm) for fascin, 56 µm (IQR = 55–57 µm) for α-actinin, and 81 µm (IQR = 70–96 µm) for CaMKII (Fig. 6 c). The apparent persistence length of the same bundles is 46 µm (IQR = 34–50 µm), 56 µm (IQR = 54–56 µm), and 16 µm (IQR = 13–30 µm), respectively (Fig. 6 d). Comparison of the fluctuation and apparent persistence lengths revealed marked differences between cross-linkers. For α-actinin bundles, the two measures were nearly identical, indicating that most bending-mode power arises from thermal fluctuations (Fig. 6 d). In contrast, fascin and especially CaMKII bundles exhibited substantially lower apparent persistence lengths than fluctuation persistence lengths, indicating that a large fraction of the bending-mode power is contained in the time-averaged bundle shape rather than in thermal fluctuations.

The fraction of bending-mode power contained in the time-averaged bundle shape can therefore be estimated as 1 – *l_p,app_*/*l_p,fluc_*, where *l_p,app_*, is the apparent persistence length and *l_p,fluc_* is the fluctuation persistence length. The fraction of bending-mode power per bundle is 2% (IQR = 1–4%) for α-actinin, 44% (IQR = 35–56%) for fascin, and 79% (IQR = 69–83%) for CaMKII. The persistence length from the cosine correlation applied to the same centerlines gives 55 µm (IQR = 39–73 µm), 84 µm (IQR = 34–101 µm), and 18 µm (IQR = 9–46 µm) (Fig. 6 e).

Raising ε from 50 to 100 kJ/mol increased the static share of the bending-mode power for fascin as well as for CaMKII, from 44% to 73% and from 79% to 83%, respectively. Over the same range, the angle between neighboring filaments in CaMKII bundles widened from 7.2 ± 0.2° to 11.0 ± 0.3° (mean ± SEM, n = 8 bundles), while its fluctuation persistence length changed by −22%. Notably, the discrepancy between fluctuation and apparent persistence lengths was greatest for CaMKII bundles, indicating that they retain a substantially larger fraction of their curvature in their time-averaged configuration than fascin- or α-actinin-cross-linked bundles.

Consistent with a conventional semiflexible polymer description, α-actinin-cross-linked bundles were well described by the persistence-length model (Fig. 6c-e). In contrast, partially fascin and mainly CaMKII bundles departed from the persistence-length fit. This departure suggests that CaMKII bundles possess structural features that are not fully captured by a simple persistence-length description at these short length scales and distinguish them from the canonical bivalent cross-linkers. This does not mean that persistence length ceases to be an effective coarse-grained descriptor of the large-scale mechanical behavior of CaMKII bundles. Rather, the departure suggests that additional structural complexity emerges at shorter length scales, where local bundle organization is not fully described by a simple persistence-length model.

One possible explanation is that the multivalent architecture and flexible linker domains of CaMKII permit local filament rearrangements or variable filament packing geometries that are not readily accessible to canonical bivalent cross-linkers. While the simulations do not directly identify the underlying structural origin of the retained curvature, they suggest that CaMKII bundles possess organizational features beyond those captured by a simple persistence-length description. Such local variations in bundle organization may contribute to the reduced persistence length and straightness ratio we measured experimentally for CaMKII-cross-linked bundles, reflected in their reduced persistence length and straightness ratio. Together, the experimental and computational results suggest that CaMKII multivalency gives rise to bundle architectures and mechanical behaviors that differ from those of canonical bivalent cross-linkers and influences actin bundle mechanics across multiple length scales.

## Conclusions

By directly comparing the morphological and mechanical properties of actin bundles formed by fascin, α-actinin, and the unique multivalent cross-linker CaMKII, we contextualized the characteristics of CaMKII bundles against those of conventional bivalent cross-linkers. We observed that the response of actin bundles to increasing cross-linker concentrations is distinct and strongly influenced by the molecular geometry of the specific cross-linker.

Consistent with previous literature (40, 64), fascin bundles stiffened significantly with increasing concentration. Notably, we observed a previously unreported ∼33% decrease in the contour length of fascin bundles at high concentrations. It remains unclear whether this shortening results from breakage due to increased stiffness and brittleness during handling, or whether it represents a self-limiting length caused by accumulated strain from fascin binding. Regardless of the specific mechanism, these findings confirm that fascin functions as a compact, rigid cross-linker, generating the straight, stiff bundles necessary for stable cellular protrusions.

In the case of α-actinin, our data showed that bundle morphology and persistence length were largely insensitive to changes in concentration. This behavior is likely due to α-actinin’s antiparallel rod geometry, which cross-links actin filaments with wide spacing (∼35 nm). This configuration creates a structure that remains sufficiently flexible such that increasing cross-linker density does not significantly alter stiffness. Consequently, α-actinin forms stable bundles with a relatively consistent mechanical baseline that may be primed for modulation by other proteins, such as myosin, during actomyosin contractions.

In contrast to both fascin and α-actinin, CaMKII bundles exhibited increased sinuosity and flexibility with increasing cross-linker concentration. We observed a decrease in the straightness ratio and a ∼33% reduction in median persistence length, a response distinct from that of the two canonical bivalent cross-linkers. Computational simulations further revealed behavior unique to CaMKII, providing evidence that its distinct mechanical response reflects differences in bundle organization rather than simply differences in bulk stiffness. Whereas α-actinin-cross-linked bundles were well described by the expected persistence-length behavior, fascin and especially CaMKII bundles departed from it. Moreover, CaMKII bundles retained a substantially larger fraction of their curvature in their time-averaged configuration than fascin- or α-actinin-cross-linked bundles, indicating that much of their bending behavior is encoded in bundle organization rather than thermal fluctuations alone.

Together, the experimental and computational results suggest that the architecture of CaMKII gives rise to structural and mechanical behavior that differs from canonical bivalent cross-linkers. One possible interpretation of these observations is the molecular structural pivot model proposed here, in which the conformational freedom provided by the flexible linker regions and multivalency of CaMKII holoenzyme permits additional organizational modes not readily accessible to bivalent cross-linkers. This interpretation is consistent with previous observations that CaMKII can support variable cross-linking angles (38, 39) and potentially facilitates inter-filament sliding (71). The increased flexibility, reduced persistence length, and enhanced retention of curvature observed for CaMKII bundles are highly compatible with the dynamic actin architectures required to support structural plasticity of small, crowded compartments such as dendritic spines. These findings provide a framework for future studies aimed at understanding how multivalent cross-linking contributes to cytoskeletal organization and structural plasticity in these cellular contexts.

More broadly, the distinct behavior of CaMKII observed in both our experimental measurements and computational simulations motivates future studies aimed at directly testing the role of multivalency in actin bundle mechanics. Engineering CaMKII variants with varying numbers of actin-binding sites, as well as performing analogous modifications to canonical cross-linkers, could help distinguish the contribution of multivalency from other structural features of the cross-linkers and provide a direct test of the ideas proposed here. A complementary approach would be to investigate how changes in holoenzyme composition influence actin bundle properties (72). When isolated from brain tissue, CaMKII exists as mixed α/β holoenzymes (73–76), and because the α subunit lacks significant actin-binding activity (20, 25), varying the α:β stoichiometry is expected to alter the effective valency and mechanical properties of the resulting actin assemblies.

Furthermore, it will be important to characterize the mechanical properties of the γ and δ CaMKII isoforms, which are also capable of cross-linking actin (25). Because multivalency and the dodecameric hub architecture are shared features of the CaMKII family, these isoforms may exhibit similar mechanical responses. Verifying this possibility could provide novel insights into the physiological roles of CaMKII-mediated actin organization in smooth muscle cells, cardiomyocytes (77–79), and other mechanically active systems (80, 81) that require a balance between structural integrity and cytoskeletal adaptability. While CaMKII is among the best characterized multivalent actin-binding proteins, it is possible to envision how assemblies of actin-cross linking proteins could produce a CaMKII-like impact on actin bundle properties. For example, α-actinin and CaMKII are both present within the post-synaptic density (66) where their association could generate highly multivalent actin-binding complexes. We hypothesize that the behaviors described here for CaMKII may extend beyond individual proteins and inform biological situations in which actin organization is governed by interconnected, multivalent binding interactions rather than simple binary interactions.

## Data availability

Data presented in this paper can be obtained from the authors upon reasonable request.

## Author contributions

S.V-H. prepared the fluorescence microscopy samples, performed the fluorescence imaging experiments, and developed the fluorescence microscopy data analysis pipeline. S.V-H., M.S., and J.P. analyzed the fluorescence microscopy data. L.L.M, C.M.D.M., and M.N.W. prepared the EM samples, performed the EM imaging experiments, and analyzed the EM data. C.B. performed and analyzed simulations, S.V-H., M.N.W., and A.-K.G. conceived the idea. P.G.W., M.N.W. and A.-K.G. supervised the research. S.V-H., C.B, M.N.W., and A.-K.G. wrote the manuscript with input from all authors.

## Acknowledgements

We thank Hugo Sanabria for valuable discussions regarding the unique properties of CaMKII. We thank Yuya Nakatani for help blinding samples, and Tyler Nelson for valuable discussions regarding statistical analysis approaches. We are grateful to Ivana Hsyung, Margareth Freire, and Diego Ledesma for their contributions to the preliminary characterization of CaMKII. We thank Matthew Swulius for his insights regarding the cryo-EM data. We also thank Nusayba El-Ali and Zoya Ahmed for their contributions to the simulation code during the Frontiers in Science (FIS) summer program at the Center for Theoretical Biological Physics. This work was supported by the National Institute of General Medical Sciences of the National Institutes of Health grant R35GM155365, the Welch Foundation grant C-2064-20210327, and startup funds from the Cancer Prevention and Research Institute of Texas grant RR200025 to A.-K.G. M.N.W. and P.G.W. acknowledge support for this work from the National Science Foundation (CHE-1743392). M.N.W. acknowledges generous support from the William Wheless III Professorship. P.G.W. was supported both by the Bullard-Welch Chair at Rice University, grant C-001, and P.G.W. and C.B. by the Center for Theoretical Biological Physics sponsored by NSF grant PHY-2019745.

