## Supplemental information for "CaMKII generates actin bundle morphology and mechanics distinct from canonical bivalent cross-linkers"

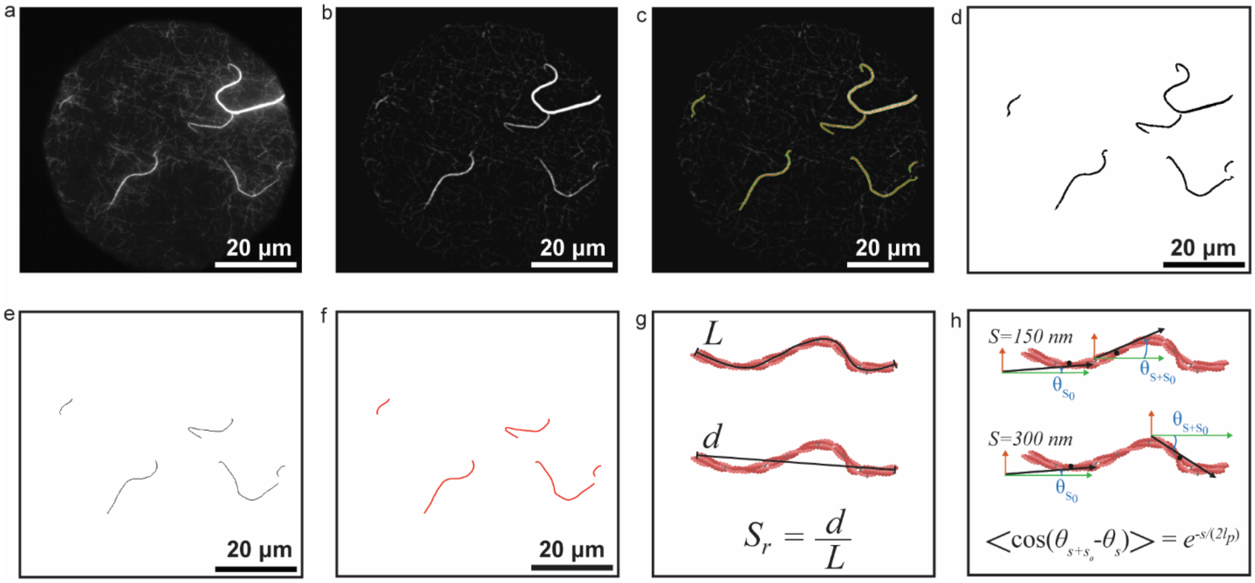


Figure S1. Imaging post-processing and analysis pipeline. (a) Summed image from a 10-frame stack. (b) Post-processed image following background subtraction, contrast enhancement, and Gaussian blur filtering. (c) Bundle detection using Ridge Detection. (d) Binary mask output generated by Ridge Detection. (e) Skeletonized image after applying a 50% intensity threshold and removing filaments extending beyond the field of view (FOV) to retain relevant bundles. (f) Smoothing of the bundle skeleton in MATLAB. (g) Schematic showing the extraction of morphological parameters: contour length (L), end-to-end distance (d), and straightness ratio ($S_r$). (h) Schematic showing the extraction of tangent angles between two segments at different separation distances s (s=150 nm and s=300 nm) to calculate persistence length ($l_p$).

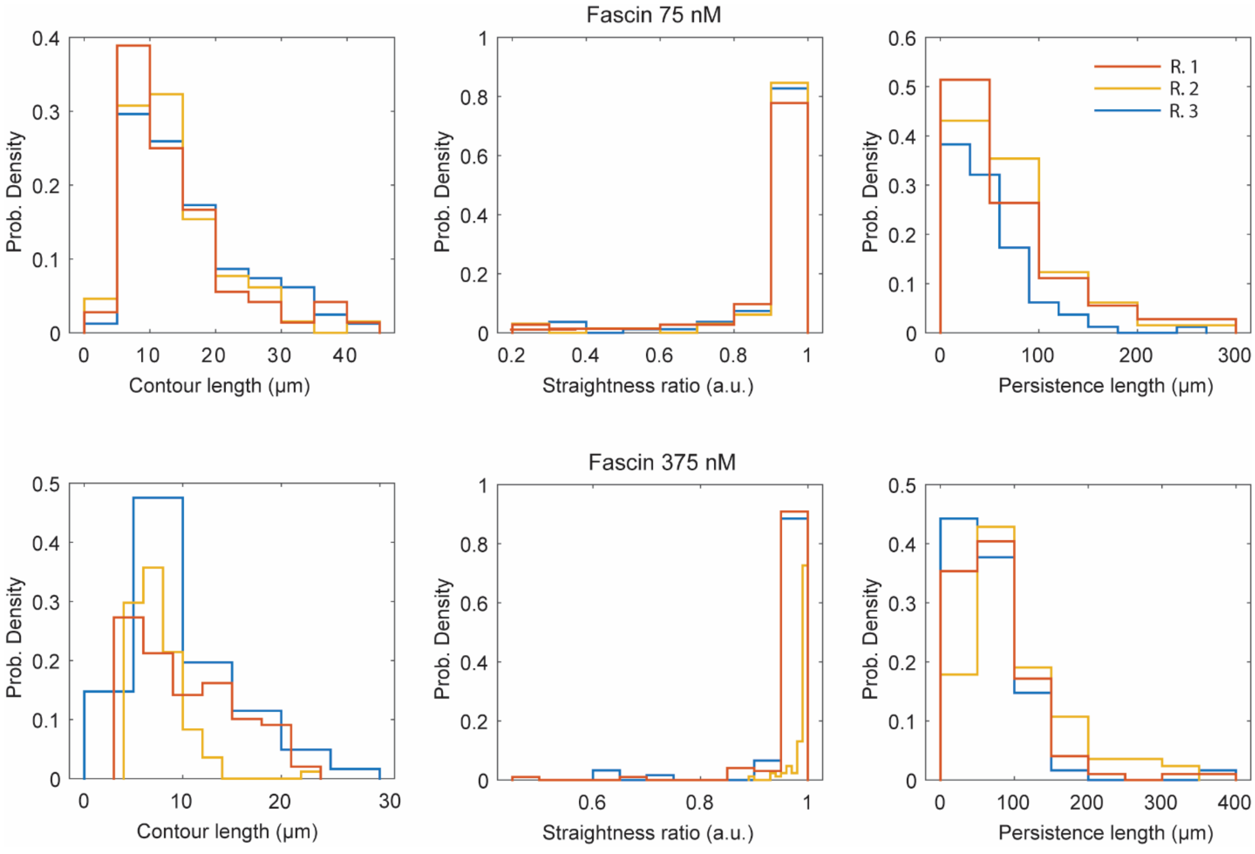


Figure S2. Distribution of parameters across replicates for fascin-cross-linked bundles. Histograms showing the probability density (Prob. Density) of contour length, straightness ratio, and persistence length for three independent replicates (R.1, R.2 and R.3) of actin bundles cross-linked by fascin at 75 nM and 375 nM.

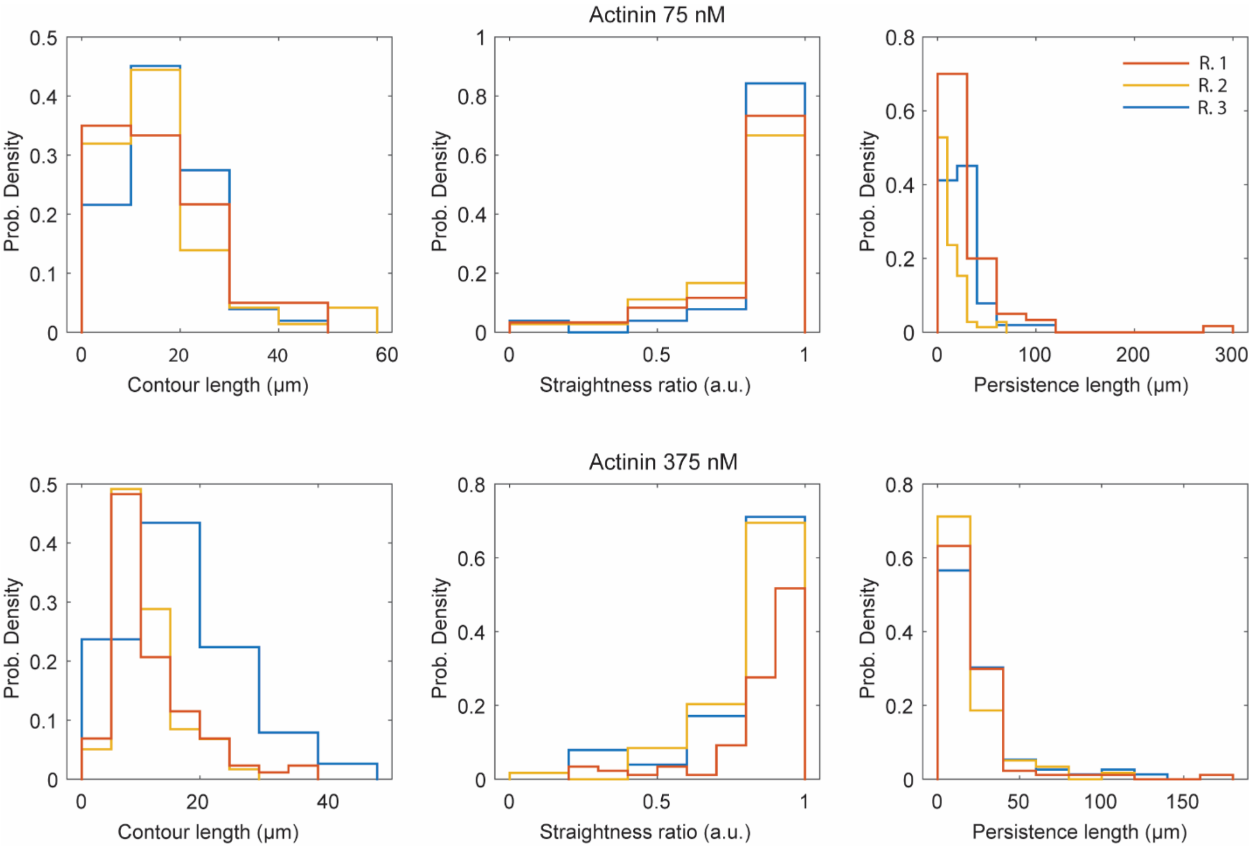


Figure S3. Distribution of parameters across replicates for α-actinin-cross-linked bundles. Histograms showing the probability density (Prob. Density) of contour length, straightness ratio, and persistence length for three independent replicates (R.1, R.2 and R.3) of actin bundles cross-linked by α-actinin at 75 nM and 375 nM.

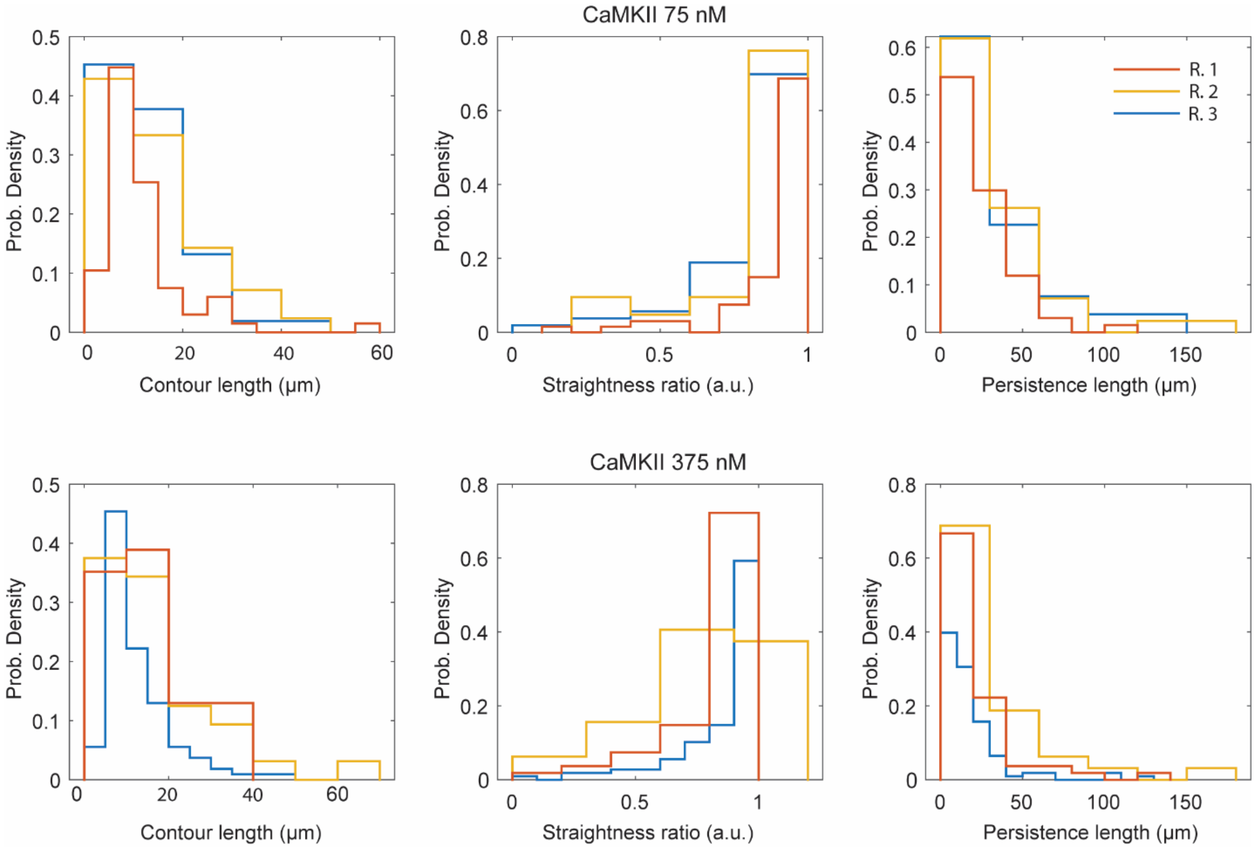


Figure S4. Distribution of parameters across replicates for CaMKII-cross-linked bundles. Histograms showing the probability density (Prob. Density) of contour length, straightness ratio, and persistence length for three independent replicates (R.1, R.2 and R.3) of actin bundles cross-linked by CaMKII at 75 nM and 375 nM.

Table S1. Assessment of persistence length fitting quality. The table displays the 95$^{th}$ percentile of the root mean square error (RMSE) for persistence length fits at each concentration (75 nM and 375 nM). This value represents the upper error threshold for 95% of the analyzed bundles. For all conditions, the RMSE remains below 0.03, indicating a good quality of fit across the dataset.

| **Cross-linker** | **95$^{th}$ Percentile RMSE** |
|---|---|
| **75 nM** | |
| Fascin | 0.0191 |
| α-actinin | 0.0275 |
| CaMKII | 0.0212 |
| **375 nM** | |
| Fascin | 0.0043 |
| α-actinin | 0.0235 |
| CaMKII | 0.0276 |

### Supplementary Methods for OpenActin simulations of actin bundles

### 1. Bundle construction

Filaments were generated by repeating one actin monomer under a screw transformation of 166 degrees and 28.2 Å per monomer, the 13/6 helix of F-actin, giving a nominal length of 1.17 µm for the 416 monomers used here.

All simulations included in the reported analysis used a minimal thin seven-filament hexagonal bundle, which although thinner than the experiments, allows us to compare the crosslinkers on the same bundle structure. Each bundle was built with the filaments straight and parallel and with the rotation of each filament about its own long axis selected by a Monte Carlo process where each filament was rotated until the connections with their neighbors was saturated.

Cross-linkers were placed by matching one binding module to the actin binding surface of each filament and accepting them when a molecule can bridge both binding modules. Two binding modules count as a candidate cross-link when the centers of mass of the putative crosslinker lie within a collision distance of each other, rejecting any that clashed with one already accepted and requiring a minimum axial separation of five monomers, 141 Å, between two cross-links bridging the same ordered pair of filaments. All valid pairs surviving that selection were kept. The resulting bundles carry 272 fascin, 256 α-actinin and 256 CaMKII molecules.

### 2. Actin model

Each actin monomer was represented by four beads, A1 to A4, corresponding to structural subdomains 1 to 4, using the ADP-state parameters of Chu and Voth (1).

In the tables below, Δi is the separation along the filament between the two monomers a term connects: 0 within one monomer, 1 nearest neighbor, 2 next-nearest. For a cross-linker, Δi = 1 denotes a bond between two binding modules of one molecule. The symbol s is reserved for arc length. Lengths are in Å and energies in kcal mol⁻¹, converted to nm and kJ mol⁻¹ at setup.

A harmonic bond is $U = \frac{1}{2} K(r - r_0)^2$ and a harmonic angle $U = \frac{1}{2} K(\theta - \theta_0)^2$. The torsion is periodic, $U = K\,[1 + cos(n\,\theta - \theta_0)]$, so $\theta_0$ is a phase while the minimum lies at $\theta_0 + 180°$.

*Table M1. Harmonic bonds*

| beads | Δi | $r_0$ (Å) | K (kcal mol⁻¹ Å⁻²) |
|---|---|---|---|
| A1–A2 | 0 | 25.85 | 4.12 |
| A1–A3 | 0 | 25.77 | 7.03 |
| A3–A4 | 0 | 24.67 | 8.64 |
| A2–A3 | 1 | 29.41 | 0.84 |
| A2–A4 | 1 | 32.65 | 0.53 |
| A4–A1 | 1 | 33.27 | 1.10 |
| A4–A3 | 1 | 31.46 | 0.40 |
| A3–A3 | 1 | 41.06 | 0.99 |
| A2–A1 | 2 | 35.18 | 0.02 |

| beads | Δi | $r_0$ (Å) | K (kcal mol⁻¹ Å⁻²) |
|---|---|---|---|
| A2–A3 | 2 | 28.61 | 0.30 |
| A4–A3 | 2 | 37.44 | 0.89 |
| A1–A1 | 2 | 57.88 | 0.67 |
| A2–A2 | 2 | 57.09 | 0.44 |
| A4–A4 | 2 | 56.93 | 0.63 |

*Table M2. Angles and torsion*

| term | beads | $\theta_0$ or phase (°) | K | K units |
|---|---|---|---|---|
| angle | A1–A3–A4 | 91.78 | 926.62 | kcal mol⁻¹ rad⁻² |
| angle | A2–A1–A3 | 94.93 | 599.12 | kcal mol⁻¹ rad⁻² |
| torsion (period 1) | A2–A1–A3–A4 | 155.89 | 477.25 | kcal mol⁻¹ |

All three terms lie within a single monomer.

*Table M3. Added flat-bottomed nearest-neighbor restraints*

In preliminary simulations the four-bead representation could reach geometries that are not representative of a native filament under sufficient thermal or mechanical stress. Therefore, we added extra energy terms to discourage those conformations. The energy is the following:

$$U(r) \ = f(x) = \begin{cases} K(r - r_{max})^2 & r > r_{max} \\ 0 & r_{min} < r < r_{max} \\ K\,(r \ - \ r_{min})^2 & r < r_{min} \end{cases}$$

Each term is zero between $r_{min}$ and $r_{max}$ and acts only outside that interval, where it limits large deformations.

| beads | Δi | r_min (Å) | r_max (Å) | K (kcal mol⁻¹ Å⁻²) |
|---|---|---|---|---|
| A2–A2 | 1 | 44.81 | 48.81 | 1.00 |
| A1–A1 | 1 | 61.36 | 64.27 | 1.00 |
| A4–A4 | 1 | 44.28 | 47.71 | 1.00 |
| A2–A4 | 1 | 39.72 | 43.53 | 1.00 |

*Table M4. Particle masses*

Assigned by the distributed force field. They are not physical molecular masses: they set the inertia of the Langevin dynamics, so the timestep that is stable and the rate at which the bending modes decorrelate. A mass of zero marks a virtual site, whose position is determined by its parents, and which carries no degrees of freedom.

| role | beads | mass (Da) |
|---|---|---|
| actin A1 | A1 | 16300 |
| actin A2 | A2 | 4060 |
| actin A3 | A3 | 11400 |

| role | beads | mass (Da) |
|---|---|---|
| actin A4 | A4 | 21500 |
| actin binding sites | Aa, Ab, Ac | 0 |
| binding module | Ca, Cb, Cd | 30000 |
| CaMKII hub | Cx1, Cx2, Cx3 | 100000 |
| CaMKII hub center and arm positions | Cc, C01–C12 | 0 |
| α-actinin rod ends | Cy1, Cy2 | 0 |

**3. Actin-binding sites**

Cross-linkers do not bind the actin beads directly. Three massless virtual sites per monomer carry the binding surface, each placed by an out-of-plane construction on the (A2, A1, A3) frame of that monomer:

$$r_{site} = r_{p1} + w_{12}\,(r_{p2} - r_{p1}) + w_{13}\,(r_{p3} - r_{p1}) + w_{cross}\,(r_{p2} - r_{p1}) \times (r_{p3} - r_{p1})$$

The site keeps a fixed position and orientation relative to its monomer, and the binding geometry is defined once rather than per cross-linker. w12 and w13 are dimensionless; the cross product has dimensions of length squared, so $w_{cross}$ has dimensions of inverse length.

*Table M5. Virtual-site weights*

| site | p1, p2, p3 | $w_{12}$(nm$^{-1}$) | $w_{13}$(nm$^{-1}$) | $w_{cross}$ (nm$^{-1}$) |
|---|---|---|---|---|
| Aa | A2, A1, A3 | 1.424234 | 0.279163 | -0.003179 |
| Ab | A2, A1, A3 | 1.116147 | 0.428060 | -0.146340 |
| Ac | A2, A1, A3 | 1.012042 | 0.186551 | -0.272158 |

Aa, Ab, Ac are the acceptor triad a binding module binds.

**4. Cross-linker models**

All three cross-linkers contain the same three-bead triangular actin-binding module (Ca, Cb, Cd), whose three sides are connected by harmonic bonds, and differ in how two or more modules are joined.

Fascin joins two modules by nine harmonic bonds, every bead of one module to every bead of the other. Its two inequivalent binding faces are taken from the cryo-EM structure of fascin cross-linking F-actin (PDB 8VO7, (2)).

α-actinin joins two modules by a single 35 nm harmonic bond between their Cb beads, representing the rod domains, with no angular or torsional restraint.

CaMKII is a dodecameric hub with twelve independent binding modules, each anchored to the hub by a bond of one consistent equilibrium length, so the dodecamer is symmetric. The hub is an equilateral triangle of three real beads Cx1, Cx2 and Cx3, carrying as virtual sites built on that triangle the hub center Cc and the twelve arm positions $Ci$, i = 1 to 12. The hub and binding geometry are taken from the structural model of the CaMKIIβ-F-actin complex of Wang *et al.* (3) assuming that the main binding site is the kinase region.

The twelve arm positions are themselves virtual sites on the hub triangle, built by the same out-of-plane construction. The hub center Cc is a three-particle average of Cx1, Cx2 and Cx3 with equal weights.

*Table M6. CaMKII hub virtual-site weights*

| arm | parents (p1, p2, p3) | w12 | w13 | $w_{cross}$ ($nm^{-1}$) |
|---|---|---|---|---|
| C01 | Cx1, Cx2, Cx3 | -0.361576 | 0.389818 | 0.054925 |
| C02 | Cx1, Cx2, Cx3 | 0.276848 | -0.305091 | 0.054925 |
| C03 | Cx2, Cx3, Cx1 | -0.361576 | 0.389818 | 0.054925 |
| C04 | Cx2, Cx3, Cx1 | 0.276848 | -0.305091 | 0.054925 |
| C05 | Cx3, Cx1, Cx2 | -0.361576 | 0.389818 | 0.054925 |
| C06 | Cx3, Cx1, Cx2 | 0.276848 | -0.305091 | 0.054925 |
| C07 | Cx1, Cx2, Cx3 | -0.305091 | 0.276848 | -0.054925 |
| C08 | Cx1, Cx2, Cx3 | 0.389818 | -0.361576 | -0.054925 |
| C09 | Cx2, Cx3, Cx1 | -0.305091 | 0.276848 | -0.054925 |
| C10 | Cx2, Cx3, Cx1 | 0.389818 | -0.361576 | -0.054925 |
| C11 | Cx3, Cx1, Cx2 | -0.305091 | 0.276848 | -0.054925 |
| C12 | Cx3, Cx1, Cx2 | 0.389818 | -0.361576 | -0.054925 |

*Table M7. Cross-linker bonds*

| role | beads | Δi | $r_0$ (Å) | K (kcal $mol^{-1}$ Å$^{-2}$) |
|---|---|---|---|---|
| binding module, all cross-linkers | Ca–Cb | 0 | 10.91 | 10.00 |
| binding module, all cross-linkers | Cb–Cd | 0 | 14.06 | 10.00 |
| binding module, all cross-linkers | Cd–Ca | 0 | 22.34 | 10.00 |
| fascin, inter-module | Ca–Ca | 1 | 73.78 | 10.00 |
| fascin, inter-module | Ca–Cb | 1 | 69.60 | 10.00 |
| fascin, inter-module | Ca–Cd | 1 | 61.55 | 10.00 |
| fascin, inter-module | Cb–Ca | 1 | 69.43 | 10.00 |
| fascin, inter-module | Cb–Cb | 1 | 65.68 | 10.00 |
| fascin, inter-module | Cb–Cd | 1 | 56.61 | 10.00 |
| fascin, inter-module | Cd–Ca | 1 | 72.16 | 10.00 |
| fascin, inter-module | Cd–Cb | 1 | 67.41 | 10.00 |
| fascin, inter-module | Cd–Cd | 1 | 56.14 | 10.00 |
| α-actinin, spectrin rod | Cb–Cb | 1 | 350.00 | 10.00 |
| CaMKII, hub triangle | Cx1–Cx2 | 0 | 49.57 | 1000.00 (10 × 100) |
| CaMKII, hub triangle | Cx1–Cx3 | 0 | 49.57 | 1000.00 (10 × 100) |
| CaMKII, hub triangle | Cx2–Cx3 | 0 | 49.57 | 1000.00 (10 × 100) |
| CaMKII, hub to arm | C0_i–Cb_i | 0 | 118.83 | 10.00 |

### 5. Repulsion interactions

$$U(r) = \left[ \varepsilon \left( \left(\frac{\sigma}{r}\right)^{12} - 2\left(\frac{\sigma}{r}\right)^{6} \right) + \varepsilon \right] \cdot \Theta(\sigma - r)$$

Non-bonded interactions energy is zero beyond σ, so each row sets a contact distance and adds no long-range attraction. σ is the sum of the two bead radii.

| beads | σ (Å) | ε (kcal mol$^{-1}$) |
|---|---|---|
| A2–A2 | 5.00 | 10.00 |
| Cc–Cc | 8.00 | 10.00 |
| A2–Cc | 6.00 | 10.00 |
| Cb–Cb | 6.00 | 10.00 |
| A2–Cb | 3.50 | 10.00 |
| Cc–Cb | 6.00 | 10.00 |
| A4–Cc | 6.00 | 10.00 |
| Cc–Ca | 6.00 | 10.00 |
| Cc–Cd | 6.00 | 10.00 |
| A4–Cb | 3.50 | 10.00 |
| Ca–Ca | 6.00 | 10.00 |
| Cd–Cd | 6.00 | 10.00 |

*Binding-site repulsion*

Binding sites on different filaments additionally repel through $U(r) = \varepsilon\,(1 - r/\sigma)^2$ for r < σ, applied per filament pair so that sites on one filament never interact. It keeps filaments from approaching closer at their binding faces than a cross-linker could bridge, and its range is below the smallest intended cross-linker spacing so that it prevents overlap without setting the lattice constant. σ = 3 nm, ε = 100 kJ mol$^{-1}$.

### 6. Binding potential

A binding module binds an actin monomer through one smooth, reversible three-point well between the module's (Ca, Cb, Cd) beads and the monomer's (Aa, Ab, Ac) virtual sites, paired one to one:

$$U = -\varepsilon_{bind} \cdot ½ \left[ e^{-\frac{\bar{d}^2}{w_1}} + e^{-\frac{\bar{d}^2}{w_2}} \right]$$

$$d^2 = \left( |Aa - Ca|^2 + |Ab - Cb|^2 + |Ac - Cd|^2 \right)/3$$

Pairing the three distances makes the well select an orientation as well as a separation. The two widths give one basin broad enough to capture an approaching module and one narrow enough to hold it in the correct conformation. The potential is continuous and bounded, so cross-links break and re-form during the simulation and the same functional form and the same widths are used for all three cross-linkers.

The well is evaluated for every module-monomer pair within the cutoff and does not by itself forbid two modules occupying one site, or one module reaching two sites at once. Occupancy is limited sterically instead: the binding module beads carry excluded volume against each other and against actin, and the binding sites of different filaments carry the additional repulsion described there, so a second module cannot approach an already occupied site closely enough to reach the narrow well.

| parameter | value | units | notes |
|---|---|---|---|
| $\varepsilon_{bind}$ | 0, 25, 50, 75, 100 | kJ mol$^{-1}$ | Binding strength |
| $w_1$ | 5 | nm$^2$ | broad capture basin |
| $w_2$ | 0.5 | nm$^2$ | narrow, orientation-selective minimum |
| cutoff | 12 | nm | |

**7. Simulation protocol**

Every condition used the settings below.

| setting | value | units |
|---|---|---|
| engine | OpenMM 8 | |
| integrator | Langevin | |
| temperature | 300 | K |
| timestep | 0.25 | ps |
| friction | 0.001 | ps$^{-1}$ |
| trajectory length | 10 | µs |
| analyzed interval | the 5 µs from 5 to 10 µs | |
| replicas | 8 | |
